# Cell-matrix force transmission regulates the loss of naïve pluripotency in mouse embryonic stem cells

**DOI:** 10.1101/2024.05.13.593501

**Authors:** Srivatsava Viswanadha, Manuel Gómez-González, Celine Labouesse, Valeria Venturini, Xavier Trepat, Kevin Chalut, Zanetta Kechagia, Pere Roca-Cusachs

## Abstract

The first step in the differentiation of mouse embryonic stem cells (mESCs) is the loss of naïve pluripotency. This step involves a major reshaping of cells from a rounded to a spread, adhesive phenotype. Whereas such reshaping is typically associated with increased force transmission to the extracellular matrix (ECM), the magnitude and role of cell-matrix forces in the loss of naïve pluripotency is unknown. Here, we show that cell-matrix forces increase during, and are required for, the loss of naïve pluripotency in mESCs. Using traction force microscopy, we show that mESCs progressively increase cell-traction forces and mechanotransduction markers as naïve pluripotency dissolves. Modulation of force transmission through myosin inhibition, substrate stiffness, or spatial differences within mESC colonies regulates the process. Increased force is triggered by GSK3-mediated signalling, and the effect of force is regulated in part by its transmission to the cell nucleus. Our work unveils a major role of cell-ECM forces in pluripotency dissolution, adding an important aspect to the interplay between biochemical and biophysical cues that drive this process.

## Introduction

How cells with identical genetic information give rise to complex self-organised multicellular organisms remains one of the most fascinating open questions in biology. This process is controlled by a series of tightly regulated biochemical and mechanical interactions. During the first stages of embryonic development, embryonic stem cells (ESCs) transition from a state of naïve pluripotency – the cellular state of the preimplantation mouse blastocyst inner cell mass – to primed pluripotency – post-implantation epiblast cells^1^. While both ESC states are pluripotent (they have the potential to give rise to cells from all three embryonic germ layers), they differ in their capacity to generate chimeras upon microinjection into blastocysts^1^. Primed pluripotency ESCs exhibit lower activity levels of pluripotency factors such as NANOG and KLF4, and increased expression of factors such as OTX2, SOX1, and Brachyury^2–4^.

Several mechanisms that drive the loss of the initial naïve pluripotency state have been described, culminating in the distinct methylation status of the key pluripotency factors and overall cell epigenetic status^5,6^. Concomitant to epigenetic differences, cells also transition from a rounded to a spread cell shape, characterised by increased cell-extracellular matrix (ECM) adhesion due to implantation^5,7–13^. This transition is further enabled by reduced membrane tension triggered by the downregulation of β-catenin signaling^8,13^. In turn, this increases endocytosis, triggering ERK signalling that drives the loss of naïve pluripotency. Beyond cell shape and tension, the process of integrin-mediated cell-ECM adhesion is typically also associated with an increase in cell-ECM force transmission^9^. This increased force transmission (i.e., traction forces) typically triggers mechanotransduction processes, which can strongly affect cell behaviour and fate^7,14^. However, cell-ECM traction forces and their potential downstream effects have not been characterised during the loss of naïve pluripotency.

In this work, we thus set out to understand the role of cell-ECM force transmission in the loss of naïve pluripotency in mouse embryonic stem cells. We found that cell-ECM traction forces increase as naïve pluripotency dissolves and that this force transmission is required for the transition. Differential force transmission can explain regional differences in the expression of key transcription factors and the dissolution of naïve pluripotency. Force transmission is triggered by signalling mediated by GSK3, and needs to reach cell nuclei to fully exert its effects.

## Results

To study the role of traction forces in the loss of naïve pluripotency, we used a well-defined and previously described cell system of mouse embryonic stem cells^1,15^. These cells can be maintained in naïve pluripotency by culturing them in N2B27 medium^16^, supplemented with two inhibitors: the MEK inhibitor, PD0325901, to inhibit ERK signalling, and CHIR99021, to inhibit GSK3. By inhibiting GSK3, CHIR99021 prevents among others β-catenin degradation, preserving β-catenin mediated signalling. This combined inhibition is known as “2i” inhibition^17^. As the 2i inhibition is removed from the cell culture medium, cells progressively lose their naïve pluripotency^17^. This process can be tracked live through a previously described Rex1-GFP fluorescent reporter mESC line^15^, which integrates a GFPd2-IRES-bsd cassette into a single allele of the Rex1 gene locus (*Zfp42*), resulting in the expression of a destabilised GFP with a half-life of 2 hours. Rex-1 expression decreases as naïve pluripotency dissolves, resulting in a progressive decrease in Rex1-GFP fluorescence^15^.

As an initial experiment, we plated mESC cells as single cells on laminin-coated polyacrylamide gels of 5kPa in stiffness (Young’s modulus), cultured in N2B27 medium supplemented with the 2i inhibitors (henceforth referred to as simply 2i medium). Cells attached, spread, and proliferated on these gels for 48 h, forming small multicellular colonies. After 48 h, we removed 2i from the medium and imaged cells over time in a confocal microscope. Removal of 2i (henceforth referred to as N2B27 medium) led to the progressive decrease in Rex1-GFP signal over 2 days (Fig. 1a,b, Sup. Video 1). In contrast, Rex1-GFP signal stayed constant over the same period in cells where 2i inhibition was maintained (Fig. 1a,b, Sup. Video 1). This process was quantified by fitting an exponential decay curve to the fluorescence curves, showing different decay constants λ between 2i and N2B27 conditions (fig. 1c). Accordingly, we also observed the expected changes in transcription factor expression that accompany the dissolution of naïve pluripotency^15,18–20^. These include an increase in OTX2 expression (Fig. S1a,b), and a decrease in the expression of NANOG (fig. S1e,f) and KLF4 (fig. S1i,j), as assessed through immunostaining carried out at different time points after 2i removal.

**Figure 1.**
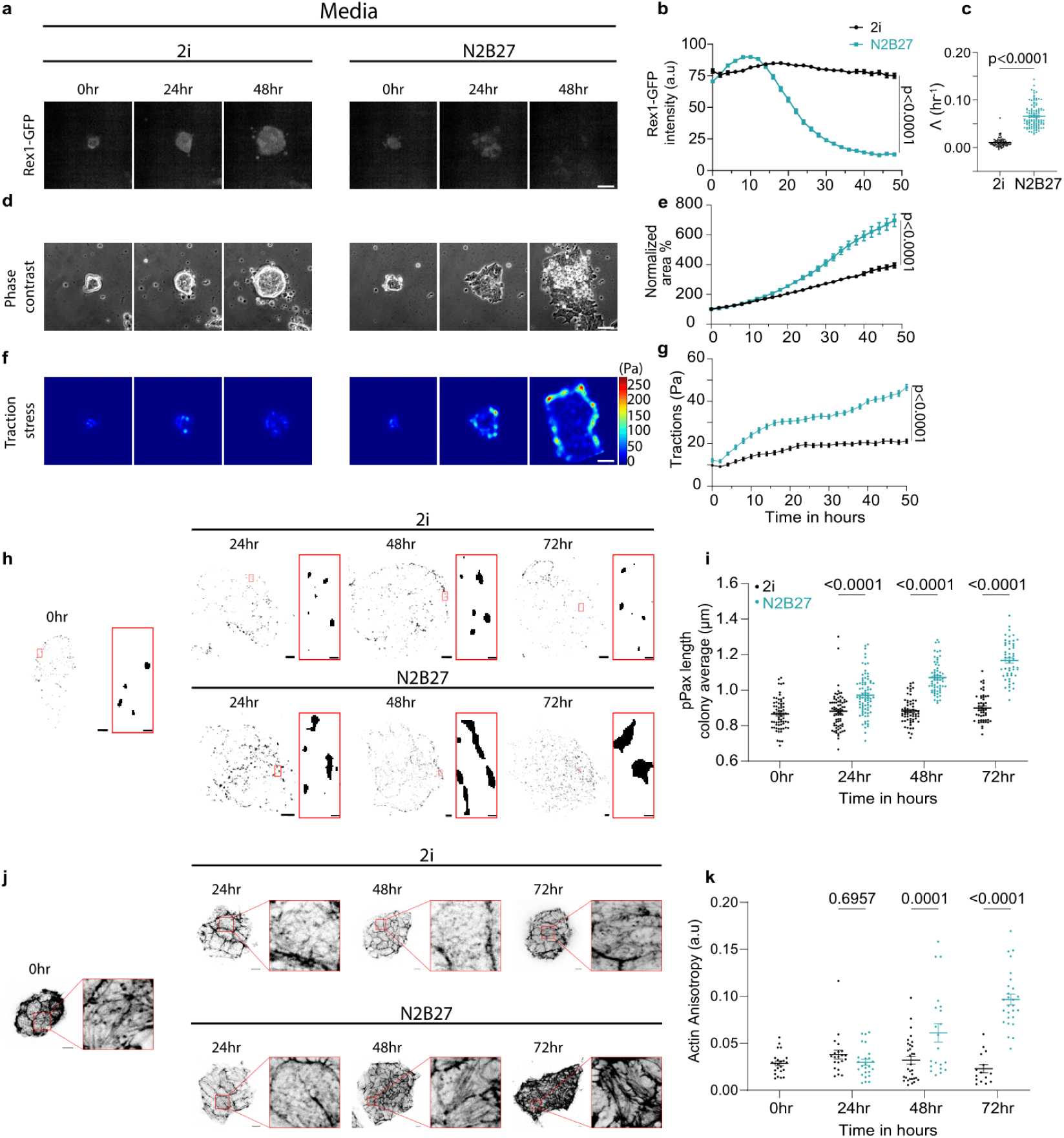
Cell-ECM forces and interactions increase as mESCs exit naïve pluripotency. a. Rex1-GFP mean intensity of mESC colonies in 2i medium (left) or N2B27 medium (right) as a function of time. The scale bar is 50 µm. b. Corresponding quantification of normalized Rex1-GFP mean intensity (3/4 independent experiments with n = 104/145 colonies for 2i/N2B27. The effect of media is significant (p<0.0001, Welsch’s t-test, for the last time point). c. Decay constants obtained from exponential fitting of Rex1-GFP decay trajectories. 3 independent experiments (n = 86/144 colonies for 2i/N2B27). The effect of media is significant (p<0.0001, Kolmogorov Smirnov test). d. Phase contrast images of mESC colonies in 2i medium (left) or N2B27 medium (right) seeded on 5 kPa polyacrylamide gels and imaged at different time points. The scale bar is 50 µm. e. Corresponding quantification of colony areas normalized to the initial value. Data from 3 independent experiments (n = 84/110 colonies for 2i/N2B27). The effect of media is significant (p<0.0001, student’s t-test, for last time point). f. Traction forces exerted by mESC colonies in 2i medium (left) or N2B27 medium (right) as a function of time. Scale bar is 50 µm. g. Corresponding quantification of traction forces averaged over colony area. Data from 3 independent experiments (n = 72/91 colonies for 2i/N2B27). The effect of media is significant (p<0.0001, Welsch’s t-test, for last time point). h. Representative images of phospho-paxillin staining as a function of time with respective zoomed insets (in red-bordered rectangles) to the right of each image. Top panel, 2i medium, bottom panel, N2B27 medium. Scale bar is 10 µm (main images) and 1µm (insets). i. Quantification of the length of phospho-paxillin adhesions. Data is from 3 independent experiments (n = 66/72/80/60/61/51/51 colonies for 2i-0hr/2i-24hr/N2B27-24hr/2i-48hr/N2B27-48hr/2i-72hr/N2B27-72hr). The effect of media and time is significant (Two-way ANOVA without repeated measures). j. Representative images of Phalloidin (F-actin), with respective zoomed areas (in red-bordered rectangles) to the right of each time point image. The top panel corresponds to 2i, and the bottom panel corresponds to N2B27 media. Scale bar is 10 µm. k. Corresponding quantification of actin anisotropy. Data is from 2 independent experiments (n = 23/21/25/31/21/15/27 colonies for 2i-0hr/2i-24hr/N2B27-24hr/2i-48hr/N2B27-48hr/2i-72hr/N2B27-72hr). The effect of media and time is significant (Two-way ANOVA without repeated measures). Error bars show mean ± s.e.m.

Exit of naïve pluripotency coincided with cell morphological and mechanical changes. First, colonies progressively grew but spread more in N2B27 than in 2i conditions (Fig. 1d,e, Sup. Video 1). Second, we measured cell-ECM traction forces using traction force microscopy as the cells transition from naïve to prime pluripotency. Average traction forces progressively increased with time but were much larger for N2B27 than 2i conditions (Fig. 1f,g, Sup. Video 1). Third, N2B27 showed a progressive increase in markers of mechanotransduction, as assessed through the growth and density of paxillin-stained focal adhesions (Fig. 1h,i, Fig. S2a) and the formation of stress fibers (Fig. 1j,k). These features were not observed in 2i samples (Fig. 1h-k).

After this initial characterization, we analysed whether cell-ECM force transmission had a causal effect in pluripotency exit, in different ways. First, we inhibited myosin contractility with blebbistatin. Force transmission was progressively impaired with increasing concentration of blebbistatin (Fig. 2a,b, Sup. Video 2). This was mirrored by the progressive slowing down of Rex1-GFP decay (Fig. 2c-e, Sup. Video 2). Of note, cell spreading was impaired in the 10 μM blebbistatin condition, possibly due to effects on cell division^21^, but not in lower concentrations, where it was even increased in a dose-dependent manner (Fig. S2b,c). This shows that cell-ECM force transmission had an effect independently of cell spreading. Second, we plated cells on substates of different stiffness. This led to small but measurable differences in forces. Consequently, we observed small differences in Rex1-GFP decay, with slightly quicker decay for stiff (15 and 5 kPa) than soft (1.5 kPa) substrates (Fig. 2f-k).

**Figure 2.**
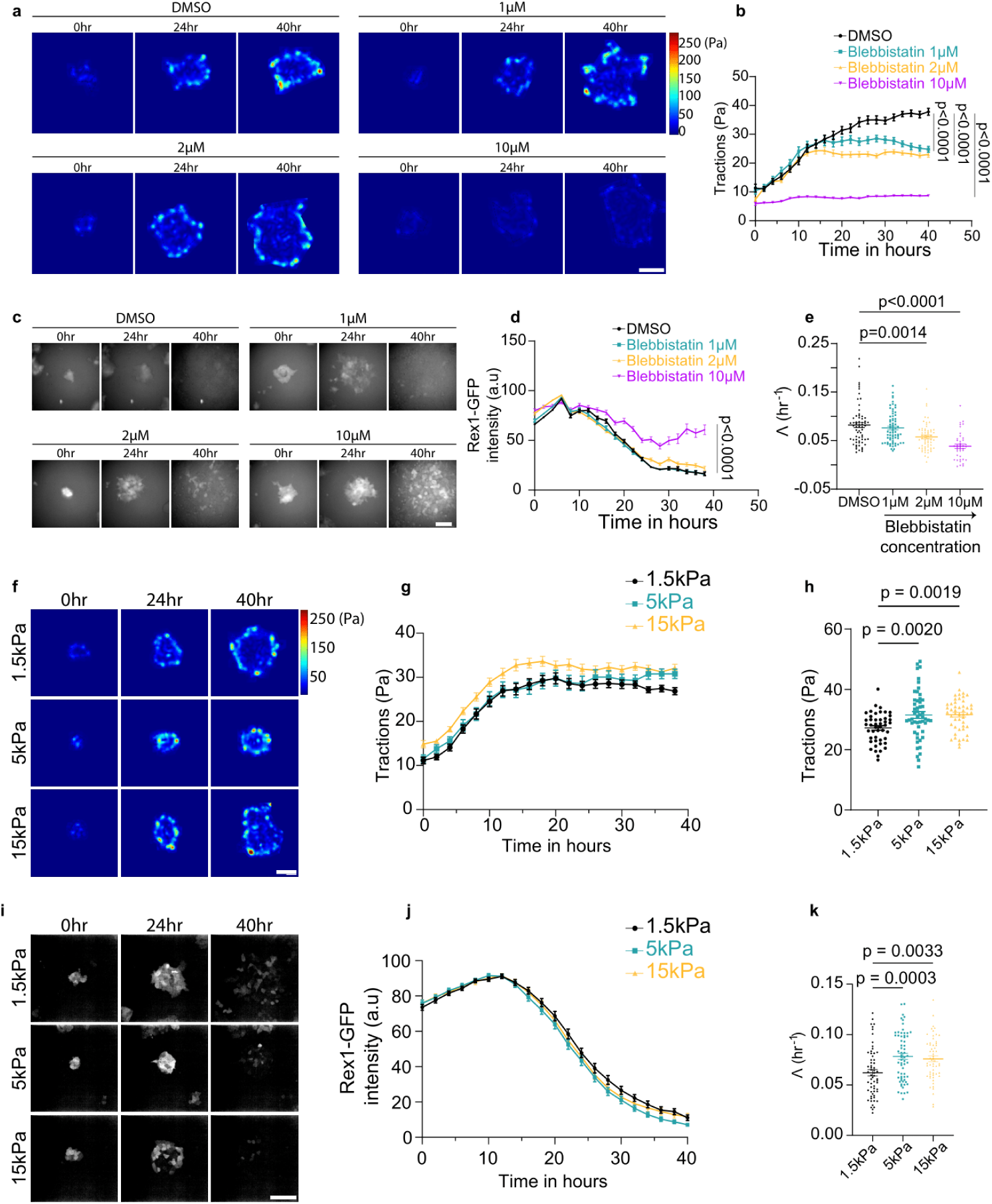
Interference with cell-ECM force transmission impairs pluripotency dissolution kinetics. a. Representative heat maps of tractions exerted by mESC colonies in DMSO or different concentrations of blebbistatin. The scale bar is 20 µm. b. Corresponding quantification of traction forces averaged over colony area. Data from 3 independent experiments (n = 58/59/40/38 for DMSO/1µM-/2µM-/10µM-Blebbistatin treatments). The effect of media is significant (p<0.0001, one-way ANOVA, for the last time point). c. Rex1-GFP signal of mESC coloni<u>e</u>s in DMSO/1µM-/2µM-/10µM-Blebbistatin treatments as a function of time. The scale bar is 40 µm. d. Corresponding quantification of Rex1-GFP mean intensity, average of 3 independent experiments (n = 65/70/61/40 for DMSO/1µM-/2µM-/10µM-Blebbistatin treatments). The effect of media is significant (p<0.0001, Kruskal-Wallis test, for the last time point). e. Decay constants obtained from exponential fitting of Rex1-GFP decay trajectories average of 3 independent experiments (n = 65/70/61/40 for DMSO/1µM-/2µM-/10µM-Blebbistatin treatments). The effect of media is significant (Kruskal-Wallis test). f. Representative heat maps of tractions exerted by mESC colonies in different stiffness conditions (Young’s modulus). The scale bar is 10 µm. g. Corresponding quantification of traction forces averaged over colony area. Data from 3 independent experiments (n = 46/52/49 for 1.5kPa/5kPa/15kPa). h. Quantification of traction force of last time point of panel g. The effect of stiffness is significant for the comparison between 1.5kPa and 15kPa, 1.5kPa and 5kPa (One-way Anova) i. Representative images of Rex1-GFP signal of mESC colonies grown on PAA gels of 1.5, 5 and 15 kPa stiffness. The scale bar is 50 µm. j. Corresponding quantification of normalized Rex1-GFP mean intensity. 3 independent experiments (n = 61/64/57 for 1.5kPa/5kPa/15kPa). k. Decay constants λ obtained from exponential fitting of Rex1-GFP decay trajectories, Average of 3 independent experiments. (n = 61/64/57 for 1.5kPa/5kPa/15kPa). The effect of stiffness is significant (One-way Anova, comparisons with respect to 1.5kPa) Error bars show mean ± s.e.m.

As a third approach to analyse the role of cell-ECM force transmission, we noticed that there were regional differences in force transmission within colonies. Indeed, cells at the edge of the colonies exerted higher forces in N2B27 conditions (Fig. 3a,b), and also exhibited much lower levels of Rex1-GFP from the beginning of the experiment (Fig. 3c-f). Cells at the edge of colonies, in fact, exhibited the same Rex1-GFP levels as single isolated cells (fig. 3c-d), suggesting that reduced cell-cell contacts explained the phenotype. Indeed, increased cell-matrix rather than cell-cell adhesion at the edges of cell colonies and monolayers has been shown to increase traction forces and mechanotransduction^22^. Consistently, mechanotransduction markers were also upregulated at the edge of the colonies, as focal adhesions were more prominent at colony edges (Fig. 1h). To assess this further, we measured the nuclear to cytoplasmic ratios of YAP, a transcriptional regulator that is a well-described marker of mechanotransduction^23–25^, and key regulator of early embryonic development, although with often contradicting roles^26–31^. Mimicking the trends of cell-matrix forces, YAP was systematically more nuclear at the edge of colonies (Fig. 3g-i). YAP nuclear levels at the colony edges also increased more consistently with time for cells in N2B27 medium, as compared to 2i (Fig. 3g-i).

**Figure 3.**
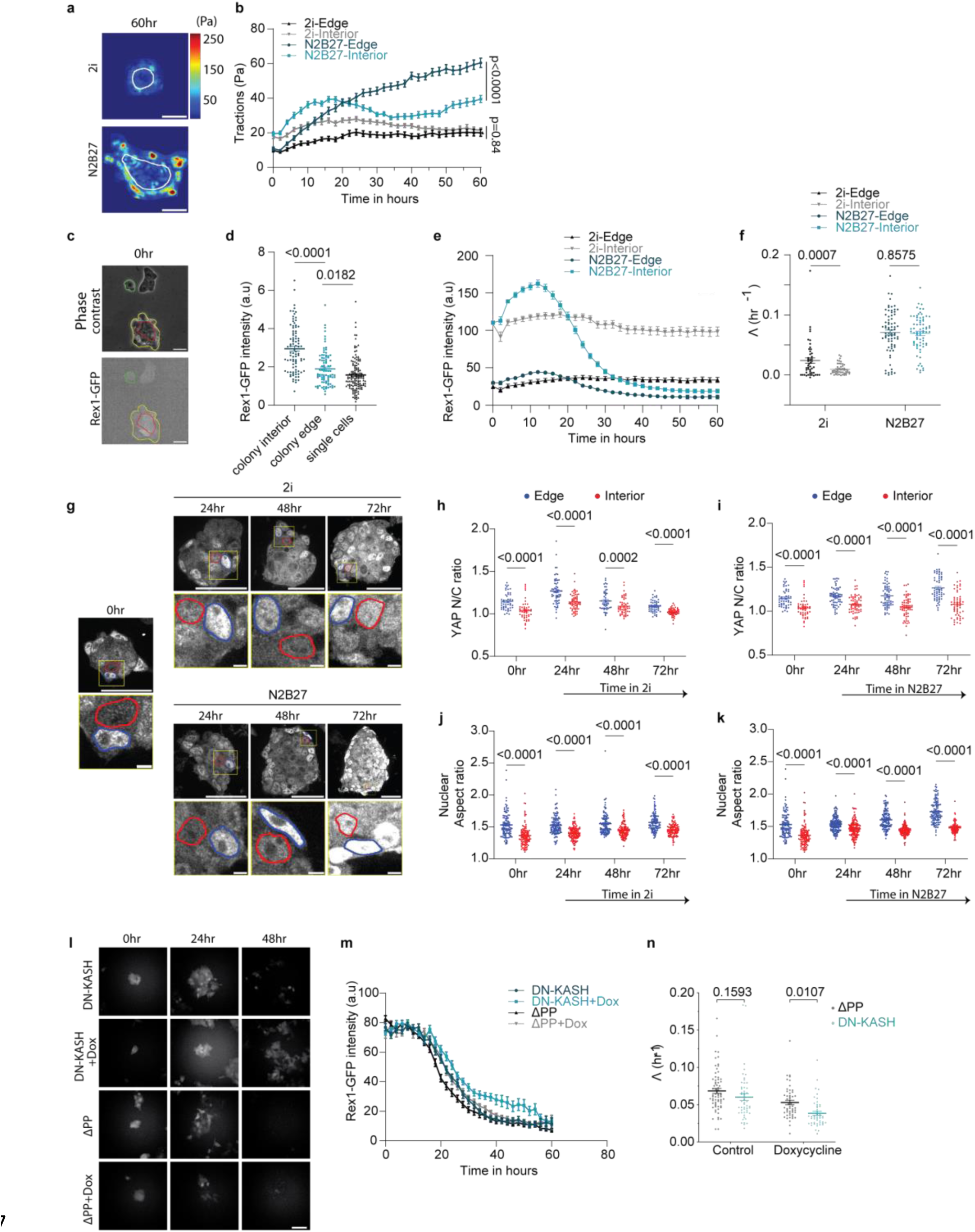
Regulation of force transmission within colonies, and to the cell nucleus, controls the kinetics of pluripotency dissolution. a. Representative traction heat maps of mESC colonies grown for 60 hours in the corresponding media. The region between within white outlines corresponds to interior, region outside white outline is edge. The scale bar is 25 µm. b. Corresponding quantification of traction forces as a function of time for edge and interior regions of mESC colonies grown in the corresponding media. Data from 3 independent experiments (n = 48/62 colonies for 2i/N2B27). The effect of Edge vs Interior is significant only for N2B27 (Two-way ANOVA with repeated measures for the last time point). c. Representative phase-contrast image (top) and Confocal fluorescence image of green channel (bottom) for mESCs immediately after media change to N2B27. The region between the yellow and red outlines corresponds to colony edge, and the region within the red outline is interior. Green outline corresponds to a single cell. Tha scale bar is 25µm. d. Comparison of Rex1-GFP mean-intensity between colony edge, colony interior and single cells, immediately after change of media to N2B27, average from 3 experiments (n = 39/39/62 colonies for Colony-edge/Colony-interior/Single-cell). Only comparison between Colony-edge and Colony-interior are significantly different (One-way ANOVA with respect to colony-edge). e. Quantification of Rex1-GFP intensity, as a function of time, resolved into edge and interior regions for mESC colonies grown in N2B27. Data from 3 independent experiments (n = 48/73 colonies for 2i/N2B27). f. Quantification of decay constants resolved into edge and interior regions for colonies grown in N2B27. Data from 3 independent experiments (n = 48/73 colonies for 2i/N2B27). The effect of Edge vs Interior is significant only for 2i. The effect of media is significant. (Two-way ANOVA with repeated measurements) g. Representative YAP fluorescence images of mESC colonies grown in 2i or N2B27 medium as a function of time, with respective zoomed images (shown in yellow-bordered rectangles). Nuclear boundaries are demarcated with outlines, blue corresponds to edge cell and red corresponds to interior cell. The scale bar is 20µm in main images and 2µm in zoomed images. h. Quantification of YAP nuclear to cytoplasmic ratio, resolved for edge and interior, and averaged for each colony grown in 2i for the corresponging time, averages from 3 independent experiments (n = 47/61/49/51 colonies for 2i-0hr/2i-24hr/2i-48hr/2i-72hr). The effect of Time and Edge vs Interior is significant (Two-way ANOVA with repeated measures). i. Quantification of YAP nuclear to cytoplasmic ratio, resolved for edge and interior, and averaged for entire colony for mESC colonies grown in N2B27 for the corresponding time, averages from 3 independent experiments (n = 47/62/62/57 colonies for 2i-0hr/N2B27-24hr/N2B27-48hr/N2B27-72hr). The effect of time and colony position is significant (Two-way ANOVA with repeated measures). j. Quantification of nuclear aspect ratios, resolved for edge and interior, and averaged for each colony grown in 2i for the corresponding time, colony averagesfrom 6 independent experiments (n = 103/127/114/104 colonies for 2i-0hr/2i-24hr/2i-48hr/2i-72hr). The effect of time and colony position is significant for all time points (Two-way ANOVA with repeated measures). k. Quantification of nuclear aspect ratios, resolved for edge and interior, and averaged for entire colony for mESC colonies grown in N2B27 for mentioned time, colony averagesfrom 6 independent experiments (n = 103/145/128/108 colonies for 2i-0hr/ N2B27-24hr/N2B27-48hr/N2B27-72hr). The effect of Time and colony position is significant for all time points (Two-way ANOVA with repeated measures). l. Representative fluorescence images of mESC colonies of DN-KASH/ΔPP genetic background grown for the corresponding time points in N2B27 on glass substrates. Doxycycline and N2B27 were added at 0hr. Scale bar is 40µm. m. Corresponding quantification of normalized Rex1-GFP intensity, as a function of time. Averaged data from 3 independent experiments (n = 50/48/70/54 colonies for DN-KASH/DN-KASH+Doxycycline/ΔPP/ΔPP+Doxycycline). n. Quantification of decay constants corresponding to Rex1-GFP fluorescence signal evolution in panel m. Data from 3 independent experiments (n = 50/48/70/54 colonies for DN-KASH/DN-KASH+Doxycycline/ΔPP/ΔPP+Doxycycline). The effect of genetic background is significant only in Doxycycline induced conditions (Two-way ANOVA). Error bars show mean ± s.e.m.

Consistent with focal adhesions and YAP trends, a similar spatial pattern was observed in key pluripotency transcription factors, indicating accelerated pluripotency exit at colony edges. Specifically, OTX2, which increases as pluripotency dissolves, showed higher expression at the edges (Fig. S1a,d). In contrast, NANOG, which decreases with the loss of pluripotency, was lower at the edges (Fig. S1e,h). Interestingly, KLF4 expression was not dependent on cell colony position but solely on 2i withdrawal (Fig. S1i,k,l), suggesting that its regulation may not depend on mechanotransduction pathways. Indeed, KLF4 is exported from the nucleus and targeted for degradation by ERK activation^32^, which is triggered as 2i is removed. Importantly, these centre/edge differences in YAP, OTX2, and NANOG were already present for cells in 2i medium, where naïve pluripotency exit was not induced (Fig. 3g,h and Fig. S1c,g). This was also the case for the Rex1-GFP signal, which showed clear centre/edge differences already at time zero upon induction (fig. 3e). This suggests that mechanical cues can affect transcriptional regulation of pluripotency exit even before biochemical induction occurs (i.e., 2i withdrawal).

Mechanical activation of YAP^24^ and other transcription factors^33^ is mediated by force transmission to the cell nucleus and subsequent nuclear deformation, which affects nucleocytoplasmic transport. Consistent with higher force transmission, nuclei showed a more elongated shape in colony edges (higher aspect ratio) and showed overall trends very similar to those of YAP (Fig. 3g,j,k). Nuclear shapes were also altered upon blebbistatin treatment, with a reduced aspect ratio (Fig. S3a,b).

The observed relationship between transcription factors, cellular forces, and nuclear shapes suggests a mechanism whereby mechanical forces are transmitted to the nucleus. To confirm this, we induced the expression of a dominant-negative KASH domain of nesprin in our cell model, following Doxycycline induction. By acting as a dominant negative, DN-KASH overexpression is known to disrupt the linker of the nucleoskeleton and cytoskeleton (LINC) complex, impairing force transmission to the nucleus^24,34^. As a control, we induced the expression of the KASH ΔPP mutant, which has impaired actin binding and does not act as a dominant negative^35,36^. We then measured the dynamics of Rex1-GFP decay upon inducing DN-KASH or KASH ΔPP. Confirming the role of cytoskeleton-nucleus force transmission, we found that DN-KASH overexpression significantly delayed Rex1-GFP decay as compared to KASH ΔPP overexpression (Fig. 3l-n). Of note, the effect was significant but rather small, likely because DN-KASH overexpression only affected nuclear force transmission and deformation mildly. In agreement with this hypothesis, changes in nuclear shapes were not detectable upon DN-KASH overexpression (Fig. S3c,d). Also, consistently, DN-KASH effects were significant on glass substrates (Fig. 3l-n) but not on hydrogels (Fig. S3e-g), where overall force transmission levels are lower, and DN-KASH effects are likely masked.

Finally, we asked how the role of cell-ECM force transmission was related to the biochemical regulation of naïve pluripotency dissolution. The main biochemical trigger of naïve pluripotency dissolution is ERK signalling, which is inhibited by PD0325901 (one of the 2i inhibitors). Additionally, dissolution also requires β-catenin phosphorylation, which is prevented by inhibiting GSK3 with CHIR99021 (the other 2i inhibitor). Inhibition of β-catenin phosphorylation with CHIR99021 stabilises cell-cell adhesions and impairs their disassembly^37,38^. To assess the respective roles of both pathways, we carried out experiments after removing only one of the two inhibitors. In these conditions, CHIR99021 inhibition clearly impaired force transmission compared to no inhibition (N2B27 medium) (fig. 4a,b, Sup. Video 3). In contrast, PD0325901 inhibition led to a traction profile similar to N2B27 medium (fig. 4a,b, Sup. Video 3). Other measurements also consistently showed stronger effects of CHIR99021 as compared to PD0325901. Rex1-GFP decay was more impaired by CHIR99021 than PD0325901 inhibition (Fig.4c-e, Sup. Video 3). Remarkably, using only CHIR99021 had decay constants similar to 2i inhibition (Fig. 4c-e, Sup. Video 3). The levels of KLF4, NANOG, and OTX2 were also observed to remain closer to 2i conditions when inhibited only with CHIR99021, than with PD0325901 (Fig. 4f-j). Finally, nuclear aspect ratios were also similar to 2i values when cells were treated with CHIR99021 rather than PD0325901, suggesting attenuated force transmission (Fig. S4a,b).

**Figure 4.**
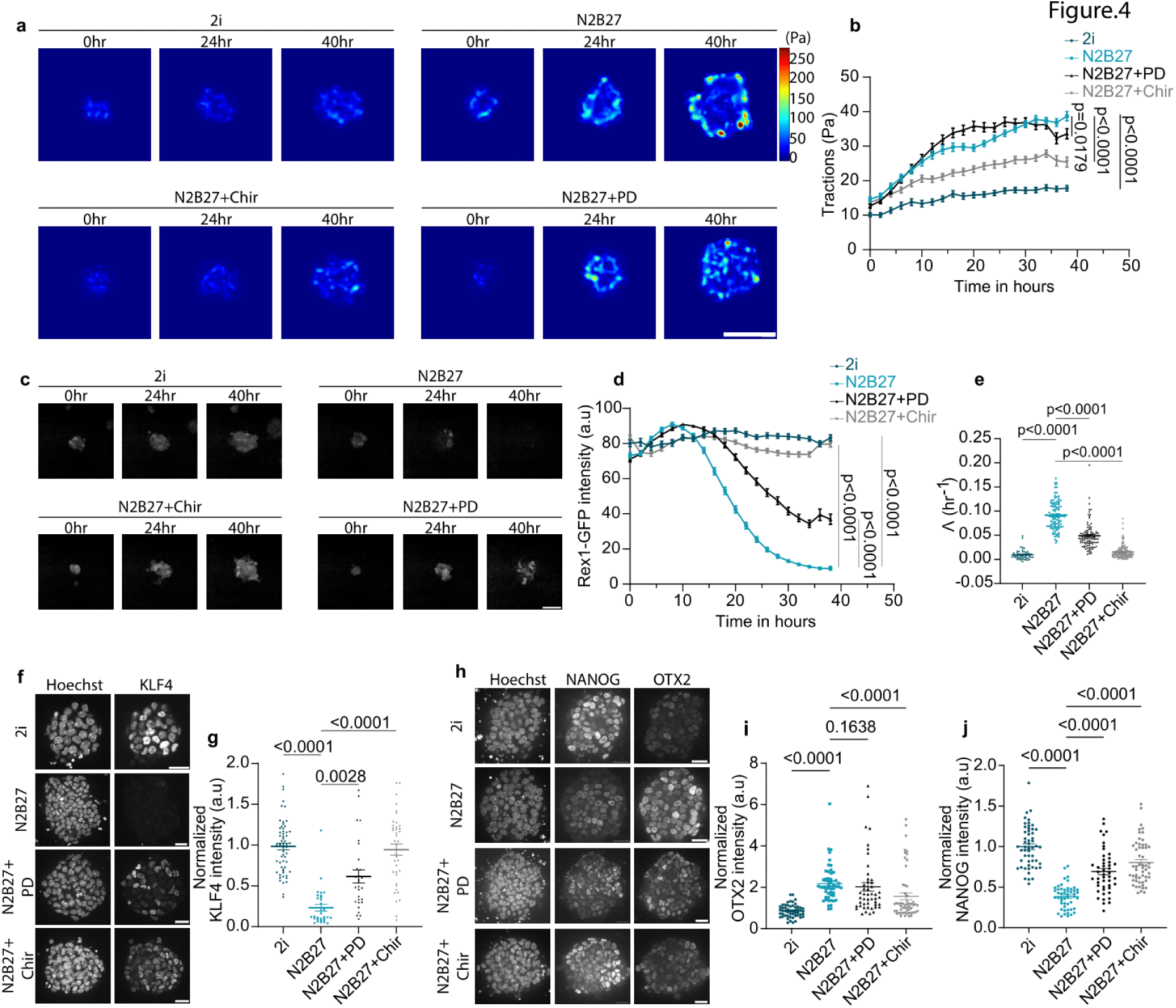
GSK3 signaling inhibition preserves naïve pluripotency more efficiently than ERK signaling inhibition. a. Representative heat maps of tractions exerted by mESC colonies in 2i/N2B27/N2B27+PD/N2B27+Chiron. The scale bar is 20 µm. b. Corresponding quantification of traction forces averaged over colony area. Averaged data from 3 independent experiments (n = 29/72/75/74 colonies for 2i/N2B27/N2B27+PD/N2B27+Chiron). The effect of media is significant (p<0.0001, Kruskal-wallis test, comparisons with N2B27, for the last time point). c. Representative images of Rex1-GFP signal of mESC colonies in 2i/N2B27/N2B27+PD/N2B27+Chiron as a function of time. The scale bar is 40 µm. d. Corresponding quantification of normalized Rex1-GFP mean intensity, averages of 3 independent experiments (n = 53/106/101/129 colonies for 2i/N2B27/N2B27+PD/N2B27+Chiron). The effect of media is significant (p<0.0001, Kruskal-wallis test, comparisons with N2B27, for last time point). e. Decay constants obtained from exponential fitting of Rex1-GFP decay trajectories from panel c, averages of 3 independent experiments (n = 53/106/101/129 colonies for 2i/N2B27/N2B27+PD/N2B27+Chiron). The effect of media is significant (Kruskal-wallis test, comparisons with N2B27). f. Representative immunofluorescence images of KLF4 after 48-hour culture on 5kPa polyacrylamide gels in 2i/N2B27/N2B27+PD/N2B27+Chiron media. The scale bar is 25µm. g. Corresponding quantification of normalized nuclear intensity averaged over the colony area, averaged data from 3 independent experiments (n = 61/31/29/40 colonies for 2i/N2B27/N2B27+PD/N2B27+Chiron). The effect of media change is significant (one-way ANOVA, without repeated measures, comparisons with N2B27). h. Representative immunofluorescence images of NANOG and OTX2 after 48-hour culture on 5kPa polyacrylamide gels in 2i/N2B27/N2B27+PD/N2B27+Chiron media. The scale bar is 25µm. i. Corresponding quantification of normalized nuclear intensity of OTX2 \, averaged data from 3 independent experiments (n =53/50/47/55 colonies for 2i/N2B27/N2B27+PD/N2B27+Chiron, one-way ANOVA, without repeated measures) j. Corresponding quantification of normalized nuclear intensity of NANOG, averaged for each colony. Data from 3 independent experiments (n =53/50/47/55 colonies for 2i/N2B27/N2B27+PD/N2B27+Chiron). The effect of media is significant Kruskal-Wallis test for OTX2, comparisons with N2B27. Error bars show mean ± s.e.m.

Altogether, these findings indicate that inhibiting GSK3 is crucial in preventing an increase in cell-matrix force transmission, and subsequent effects in naïve pluripotency dissolution.

## Discussion

In this work, we demonstrate that cell-ECM force transmission gradually increases as pluripotency is dissolved and is necessary to exit naïve pluripotency. Force transmission is enabled by GSK3 activation. Whereas other roles of GSK3 cannot be discarded, its effects in naïve pluripotency dissolution are likely mediated by its inhibition of β-catenin phosphorylation and degradation, as shown previously^8,13^. Mechanically, due to β-catenin’s role in cell-cell contacts, its degradation may shift cell adhesion from cell-cell to cell-matrix adhesions, promoting cell-matrix force transmission. This hypothesis is supported by our finding that pluripotency dissolution is favoured at colony edges, where matrix interactions and traction forces are higher. Notably, even under 2i conditions, signs of partial pluripotency dissolution (as indicated by higher OTX2 and nuclear YAP levels, and lower Rex1-GFP signal) are evident in isolated cells and those at colony edges. Our findings also implicate nuclear force transmission as a mediator, potentially by influencing transcription factor localisation and function^24,39,40^.

Previous work carried out before the introduction of the 2i inhibition system (and therefore lacking the naïve pluripotency signature) showed that stiff substrates promote later stages of mESC differentiation, and that the expression of related pluripotency factors (such as NANOG and SOX2) depend on the LINC complex^9,41^. Our findings show that this mechanical effect is triggered as early as the dissolution of naïve pluripotency, and that it is mediated by cell-ECM force transmission. More recently, reduction in β-catenin signaling in naïve mESCs was shown to decrease RhoA activity, decreasing membrane tension, allowing cell spreading, and enabling ERK signaling and subsequent naïve pluripotency dissolution through endocytosis^8,38^. Thus, membrane-tension induced cell spreading most likely acts upstream not only of ERK signalling, but also of cell contractility, with both having an important role. Interestingly, ERK inhibition alone did not greatly impair the dynamics of pluripotency dissolution (Fig. 4c-e). This result, combined with the decrease in in Rex1-GFP signal in cells at colony edges even in 2i conditions, suggests that mechanical forces can at least partly compensate for the role of ERK^8,38^

The potential mechanisms by which force transmission from the matrix to cells and their nuclei regulate naïve pluripotency dissolution remain unclear. In this regard, our previous research has established that force transmission to the nucleus regulates the nuclear translocation rates of various transcription factors and proteins^24,33^. The potential involvement of this mechanism in regulating transcription factor levels in mESCs warrants further investigation. Notably, OTX2, a transcription factor associated with early differentiation, exhibits a pronounced distribution gradient from the edge to the interior of cell colonies, independent of biochemical cues, contrary to NANOG distribution. This finding complements existing research suggesting an antagonistic relationship between OTX2 and NANOG ^42^, and highlights the multiple layers of regulation and complex interactions among transcriptional regulators.

Similarly, the function of YAP in mouse embryonic stem cell (mESC) differentiation is controversial; it can act as both a facilitator and an inhibitor of this process ^30,31,43^. Our findings suggest that the spatial variation in nuclear YAP, specifically its increased nuclear accumulation at the colony edge, may underlie the observed acceleration in differentiation. Indeed, this gradient of YAP activity could be a decisive factor for cell fate transitions, particularly as low YAP levels are associated with gene expression and high levels with gene suppression of key pluripotency factors such as OCT4 and NANOG^43^ .

The influence of mechanical forces on gene regulation is well-established across various contexts^30,44–46^. Particularly during the early stages of embryonic development, cells are distinguished by remarkably pliable nuclei due to low Lamin A/C levels and a state of open chromatin ^47–51^. Low nuclear mechanical resistance may render embryonic nuclei sensitive to even minor mechanical cues, which could precipitate changes in gene expression. While it remains to be conclusively determined whether mechanical forces are the primary drivers of symmetry breaking and differentiation in this context, our findings suggest this possibility. In any case, our results demonstrate a major role of mechanical force in the earliest step of cell differentiation, with likely implications in multiple scenarios in development.

## Methods

### Cell culture

Mouse embryonic stem cells (mESCs) from the previously characterized Rex1-GFP line^52^ were cultured on tissue culture plastic Nunc plates that had been coated with 10µg/ml Laminin-111 (Sigma-Aldrich L2020) in PBS, and incubated for at least 2 hours at 37°C. Cells were cultured in N2B27 medium, with the addition of 100 U/ml LIF (Merck Millipore ESG1106), 3 µM Chiron or CHIR99021 (Sigma-Aldrich SML1046), and 1 µM PD0325901 (Sigma-Aldrich PZ0162). The combination of CHIR99021and PD0325901 inhibitors is referred to as “2i” throughout the manuscript. Cells were passaged on alternate days using Accutase (Millipore SCR005) and regularly tested for mycoplasma. Media was changed every day. Cells were cultured without LIF for the passage before experiments.

N2B27 defined basal medium contained 1:1 Neurobasal and DMEM/F-12 medium (Sigma-Aldrich D6421), Neurobasal medium (Life technologies 21103-049), 2.2 mM L-Glutamine (Thermo Fisher – 25030081), 0.5x B27 (Invitrogen – 17504044), 3 µM CHIR99021 (Sigma-Aldrich SML1046), 1 µM PD0325901 (Sigma-Aldrich PZ0162), 50 mM β-Mercaptoethanol, 12.5 ng.mL^-1^ Insulin zinc (Sigma-Aldrich I9278), 0.1% Sodium bicarbonate (Thermo Fisher – 25080094) and 0.5% v/v 200x N2.

N2 was prepared in-house. 200x N2 was made using 8.791 mg.mL^-1^ Apotransferrin (Sigma-Aldrich T1147), 1.688 mg.mL^-1^ Putrescine (Sigma-Aldrich P5780-5G), 3 μM Sodium Selenite (Sigma-Aldrich S5261), and 2.08 µg.mL^-1^ Progesterone (Sigma-Aldrich P8783-1G). All the above components were dissolved in DMEM/F-12 (Sigma-Aldrich D6421).

For passaging, cells were treated with accutase for 5 minutes at room temperature. Followed by neutralization with 8 times its volume with BSA fraction V (1.5% v/v in DMEM/F-12). Cells were seeded at a density of 15,000 cells.cm^-2^ on Laminin-111 coated Nunc plates.

For experiments, cells were seeded after passage as single cells on laminin-coated coverslips or polyacrylamide gels (see below) at a density of 15000 cells.cm^-2^ in N2B27 +2i medium. After 48 h, cells were washed with N2B27, and treated with the corresponding medium (N2B27 medium with/without the 2i inhibitors) or drugs (blebbistatin or doxycycline). This timepoint was considered to be the start of experiments, and image acquisition (for timelapse experiments) started immediately therafter.

### Blebbistatin treatment

The para-nitro derivative of blebbistatin was chosen for myosin inhibition experiments due to its non-phototoxic and non-cytotoxic effects. The drug was purchased as lyophilized powder (Motorpharma, cat.no. 1621326-32-6) and dissolved in 100% DMSO (Sigma-Aldrich, cat. no. D8418) to prepare a stock solution of 10mM concentration that is stored at -20^0^ C. For experiments involving blebbistatin treatment, para-nitro blebbistatin was added to N2B27 media to arrive at final concentrations of 1, 2 and 10μM. As a control, DMSO was added to the media at the same volumetric proportions used for 10μM para-nitro blebbistatin.

### DN-KASH experiments

Retroviral particles for the generation of DN-KASH and Delta-PP lines were generated in HEK293T cells expressing retroviral packaging plasmids (gift from N. Montserrat) and transfected using Lipofectamine™ 3000 Transfection Reagent (Invitrogen). The viral titer collected 2-days post transfection was used to infect Rex1-GFP mESCs overnight. Rex1-GFP cells with stable integration of viral DNA was selected through 400µg/ml Geneticin (Fischer scientific, cat.no. 10204773) treatment for 6 days. Later, selection for high inducible expression was done using FACS after subjecting the cells for 1µg/ml doxycycline treatment for 16 hours. Cells were maintained in 2i thereafter.

### Preparation of polyacrylamide gels

Cover glasses of No.1 thickness (Superior Marienfeld) were used as top coverslips and were treated with Surfasil (20% v/v in Chloroform) for 2 hours at room temperature followed by quick wash with 100% methanol and air-drying. Glass bottom MatTek dishes were activated with a solution of acetic acid, 3-(Trimethoxysilyl) propyl methacrylate (Sigma), and 96% ethanol (1:1:14 in volume ratio) for 20 min at room temperature. The dishes were then quickly washed twice with 96% ethanol and air-dried. Different concentrations of acrylamide and bis-acrylamide were mixed to produce gels of different rigidity^53^ and mixed with 0.04% v/v fluorescent carboxylated 200 nm beads (Invitrogen), 0.05% APS (Sigma-Aldrich A3678), and 0.05% v/v tetramethylethylenedi-amine (Sigma-Aldrich T9281). Specifically, concentrations of acrylamide/bis-acrylamide (in %, w/v) were of 5/0.04, 7.46/0.044, and 7.5/0.16 for gels of 1.5. 5, and 15 kPa in Young’s modulus respectively (as calibrated previously^25,54^). The solution was placed on the silanized glass surface and covered with surfasil treated coverslip, letting the gel to polymerize for 50 min. The coverslip was then removed, and gels were washed twice with 1X-PBS. Gels were then coated with Sulfo-SANPAH by applying 2mg/ml solution and exposing to UV light (15 watt and 365nm wavelength) for 7.5 minutes followed by two washes with 10mM Hepes and one wash with 1X-PBS. Finally, gels were incubated overnight at 4° C with 100 µg/ml Laminin-111 (Sigma-Aldrich L2020) in 1X-PBS. The rigidity of polyacrylamide gels was measured and characterised with Atomic Force Microscopy as described previously^53,55^. For immunofluorescence samples, non-fluorescent carboxylated latex beads were used during gel preparation.

Before experiments, the protein-coated gels were sterilized with 15-minute UV treatment and equilibrated with 2i media at 37°C for 30 minutes. mESCs were seeded as single cells at 12,000 cells.cm^-2^. Immediately before the experiments, cells were washed with N2B27 media.

### Traction force microscopy

For traction force experiments, cells were seeded on PAA gels of 5 kPa unless specified (Fig. 2-j). Traction force experiments were carried out using multidimensional acquisition routines on an automated inverted microscope (Nikon Eclipse TiE) with a spinning disk confocal unit (CSU-WD, Yokogawa) and a Zyla sCMOS camera (Andor). Micromanager software was used, with a custom-made script allowing beads channel to be imaged with a z-step of 0.5 μm, and the phase-contrast and GFP channels to be imaged with a z-step of 2 μm. The blebbistatin TFM experiments were performed using an automated inverted microscope (Nikon Eclipse Ti) with a MetaMorph/NIS Elements imaging software. In all cases, a 40x objective (S-plan fluor, Ph2; NA, 0.75) was used. Both microscopes were equipped with a temperature box maintaining 37 ^0^C and a chamber maintaining CO2 and humidity.

During time lapses, we acquired images of the Rex-GFP signal (green), bead fluorescence (red), and phase contrast images of the cells. Images were acquired every 2 hours during the experiment. A reference image of beads was obtained at the end of experiments, upon cell trypsinization. Gel deformation maps between the different experimental time points and the reference image were computed with a home-made particle imaging velocimetry (PIV) software^56^ in Matlab (MathWorks Inc.) as described previously^56^. Traction forces were computed from deformation maps using Fourier traction microscopy with a finite gel thickness as described previously^57^, and averaged for each cell.

### Immunofluorescence

Cells were fixed with 4% formaldehyde for 10 minutes at room temperature followed by 3 washes with 1X PBS. Cells were then permeabilized with 0.1% Triton-X (in PBS) for 20 minutes followed by a 10-minute wash with 2% w/v BSA (in PBS). Blocking was performed with 2% w/v BSA + 2% v/v FBS in 1X-PBS for 30 minutes at room temperature. Cells were then incubated overnight at 4°C with the primary antibody. Secondary antibodies and Phalloidin-atto 488 (Sigma-Aldrich, Cat# 49409, when used) were added for 2 hours at dilutions 1:300 and 1:1000, respectively. Hoechst 33342 (Invitrogen, Cat# H3570) was used to label nuclei at 2µg/ml for 20 minutes at room temperature in blocking buffer. Finally, cells were washed with blocking buffer 4 times (1-quick and three 10 minute) and stored in 1X/PBS. All the samples were imaged the following day.

The primary antibodies used, and their respective dilutions are: Rabbit Phospho-Paxillin 1:100 (Tyr118) (Cell Signaling, Cat# 69363), mouse anti-YAP1 (63.7) 1:100 (Santa Cruz, Cat# sc-101199), rat Nanog 1:200 (Thermo Fischer eBioMLC-51, Cat# 14-5761-80), goat KLF4 1:400 (R&D, Cat# AF3158), goat Otx2 1:300 (R&D, Cat# AF1979), rabbit Sox1 1:200 (Cell Signaling, Cat#4194).

The secondary antibodies used are: goat anti-mouse Alexa Fluor − 488 (Cat# A-11029), −555 (Cat# A-21424), −647(Cat# A-21236), goat anti-rabbit Alexa Fluor −555 (Cat# A-21429), donkey anti Rabbit −488 (Cat# A-21206), −647 (Cat# A-31573), donkey anti-rat Alexa Fluor −488 (Cat# A-11006), −555 (Cat# A-21434), −647(Cat# A-21247), donkey anti-goat Alexa-Fluor −488(Cat# A-11055), -555(Cat# A-32816),−647(Cat# A-21447), and donkey anti-mouse Alexa-Flour −488(Cat# A-21202), −647(Cat# A-31571) all at 1:300 concentration (ThermoFisher).

### Image acquisition

Immunofluorescence images were taken in a Nikon TiE inverted microscope with a spinning disk confocal unit (CSU-WD, Yokogawa) and a Zyla sCMOS camera (Andor), using 60x objective (plan apo; NA, 1.2; water immersion). Time-lapse epifluorescence images were taken on an automated inverted microscope (Nikon Eclipse Ti) using MetaMorph/NIS Elements imaging software and 40x objective (S-plan fluor, Ph2; NA, 0.75). Time lapse confocal images were taken in a Nikon TiE inverted microscope with a spinning disk confocal unit (CSU-WD, Yokogawa) and a Zyla sCMOS camera (Andor), using 60x objective (S-plan fluor, Ph2; NA, 0.75).

### Image analysis

Rex1-GFP images were obtained from confocal images as explained above. To quantify images, the mean intensity over the colony area was calculated. To calculate decay constants λ, Rex-GFP quantifications over time for each colony were fitted from the point of maximum intensity with an exponentially decaying function as: (𝑁𝑜𝑟𝑚𝑎𝑙𝑖𝑧𝑒𝑑 𝑚𝑒𝑎𝑛 𝑖𝑛𝑡𝑒𝑛𝑠𝑖𝑡𝑦) = 100𝑒^−𝜆𝑡^.

For immunostainings, Fiji software was used to perform the image analysis. The length of p-Paxillin focal adhesions was assessed using the basal plane of confocal images by measuring the length of bright focal adhesions (FA) and averaging the length of all FAs per colony. YAP n/c ratios were calculated using the basal plane of confocal images and by dividing the intensity of a nuclear region and a region with equal size in the cytosol immediately adjacent to the nuclear region, upon correcting for background. The corresponding Hoechst staining image and fluorescent staining signals were used to delimit nuclear versus cytosolic regions. For transcription factor expression, the areal mean intensity of the fluorescence within the nuclear region after background correction is measured and averaged for all nuclei per colony. Values were normalized to the 2i condition at 0 hours of the corresponding experiment.

For measurements of nuclear aspect ratio, the Hoechst channel was used to manually delimit the nuclear region and an ellipse was fit to the ROI using the ImageJ in built function. The ratio of major axis to minor axis was calculated and considered as the aspect ratio of the given nucleus. The signal corresponding to a colony was obtained by averaging all the nuclei aspect ratios. Only basal plane nuclei are considered for this analysis.

Actin anisotropy was quantified in basal planes of cell colonies labelled with phalloidin. Each value represents an average of 3 cells per colony. The anisotropy quantification was implemented using the ImageJ FibrilTool plug-in^58^.

### Resolving Edge and Interior regions

For time lapse imaging, using the colony outlines drawn on the phase contrast images, a concentric region eroded through eucledian distance map to 80% of the original ROI radius was obtained. The region contained within it is labelled interior and the region in the remaining 20% of the rim is termed edge. The areal mean of traction stresses within these Interior and Edge regions was calculated from the traction maps.

For immunofluorescence images, the outer most cells of the colony define the edge region. The rest of the colony is considered the interior region. The mean intensities of all edge/interior cells are averaged and taken as the edge/interior signal of the particular colony. The edge and the interior signals for a given colony are normalized to the average of mean intensities of all colonies pertaining to the 2i-0hr condition (control).

### Statistical analysis

Statistical analysis was performed using GraphPad Prism software (GraphPad, version 9). Statistical significance was determined by the specific tests indicated in the corresponding figure legends. Non-parametric tests were performed when both original and log-10 transformed datasets were not normally distributed where appropriate. All experiments presented in the manuscript were repeated in at least 2 independent experiments.

## Author contributions

Conceptualization: S.V, Z.K, P.R.-C.

Resources: C.L., K.C.

Data curation: S.V., Z.K., P.R.-C.

Software: M.G.

Formal analysis: S.V., Z.K.

Supervision: Z.K., P.R.-C.

Funding acquisition: X.T., K.C., P.R.-C.

Validation: Z.K., P.R.-C.

Investigation: S.V., Z.K.

Visualization: S.V., Z.K.

Methodology: S.V., M.G., C.L., V.V., X.T., K.C., Z.K.

Writing—original draft: S.V., Z.K., P.R.-C.

Project administration: P.R.-C.

Writing—review and editing: S.V., Z.K., P.R.-C.

## Data availability

Source data for all figures is available as a supplementary file.

## Acknowledgements

We thank M. Purciolas-Casas, M. Morcillo, and E. Coderch for providing technical support. We acknowledge funding from “la Caixa” Foundation (grant LCF/PR/HR20/52400004 to X.T., and P.R.-C.), the Spanish Ministry of Science and Innovation (PID2022-142672NB-I00 to P.R.-C., PID2021-128635NB-I00 to X.T.), the Generalitat de Catalunya (2021 SGR 01425 to X.T. and P.R.-C.), The prize “ICREA Academia” for excellence in research to P.R.-C., Fundació la Marató de TV3 (201936-30-31 to P.R.-C.), and the European Research Council (grant 101097753 MechanoSynth to P.R-C. and Adv-883739 Epifold to X.T.). IBEC is a recipient of a Severo Ochoa Award of Excellence from MINCIN.

