## Supplemental Information for "Cell-matrix force transmission regulates the loss of naïve pluripotency in mouse embryonic stem cells"

### Supplementary figures

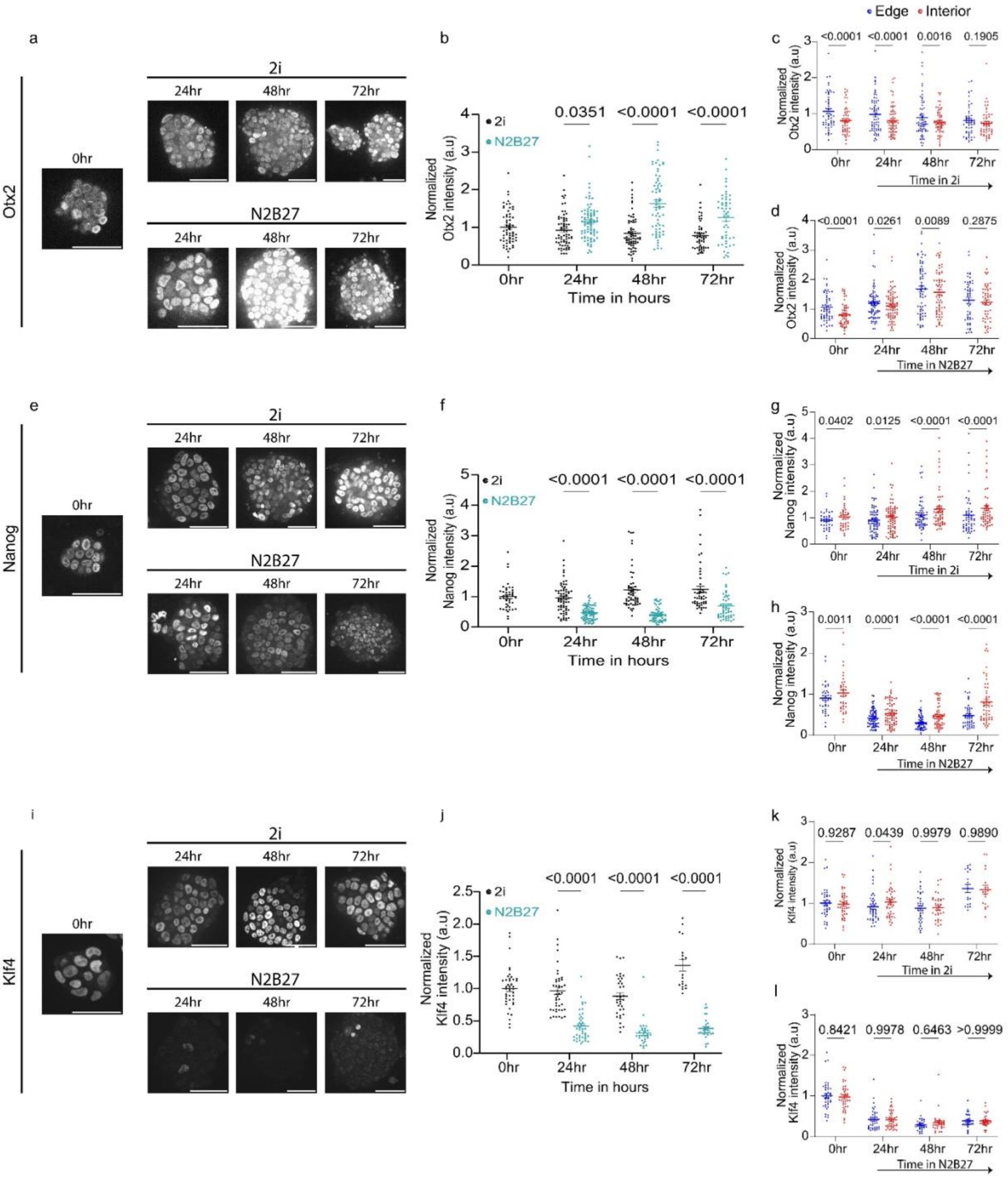


**Supplementary Figure 1. Characterization of pluripotency transcription factors.**

a. Representative images of mESC colonies immunostained for OTX2 in 2i (top)/N2B27 (bottom). The scale bar is 50 µm.

b. Corresponding quantification of normalized OTX2 nuclear intensity (normalized to average of 2i in corresponding experiment). Data from 3 independent experiments (n = 62/67/82/65/66/54/52 colonies for 2i-0hr/2i-24hr/N2B27-24hr/2i-48hr/N2B27-48hr/2i-72hr/N2B27-72hr). The effect of media change and time is significant (Two-way ANOVA).

c. Corresponding quantification of OTX2 nuclear intensity in 2i, colony edge vs Interior. Averages of 3 independent experiments (n = 57/67/65/54 colonies for 2i-0hr/2i-24hr/2i-48hr/2i-72hr).

The effect of edge vs interior is significant for all except 2i-72hr, the effect of time is not significant (Two-way ANOVA with repeated measures).

d. Corresponding quantification of OTX2 nuclear intensity in N2B27, colony edge vs interior. Averages of 3 independent experiments (n = 57/83/66/52 colonies for 2i-0hr/N2B27-24hr/N2B27-48hr/N2B27-72hr). The effect of edge vs interior and of time is significant for all except N2B27-72hr (Two-way ANOVA with repeated measures).
e. Representative images of mESC colonies immunostained for Nanog in 2i (top)/N2B27 (bottom). The scale bar is 50 μm.
f. Corresponding quantification of normalized NANOG nuclear intensity. Data from 3 independent experiments (n = 43/69/74/55/60/52/52 colonies for 2i-0hr/2i-24hr/N2B27-24hr/2i-48hr/N2B27-48hr/2i-72hr/N2B27-72hr). The effect of media and time is significant (Two-way ANOVA).
g. Corresponding quantification of NANOG nuclear intensity in 2i, colony edge vs interior. 3 independent experiments. (n = 38/69/55/52 colonies for 2i-0hr/2i-24hr/2i-48hr/2i-72hr) The effect of edge vs interior and time is significant (Two-way ANOVA with repeated measures).
h. Corresponding quantification of NANOG nuclear in N2B27, colony edge vs interior. Averages of 3 independent experiments (n = 38/74/60/52 colonies for 2i-0hr/N2B27-24hr/N2B27-48hr/N2B27-72hr). The effect of edge vs interior and time is significant (Two-way ANOVA with repeated measures).
i. Representative images of mESC colonies immunostained for Klf4 in 2i (top)/ N2B27 (bottom). The scale bar is 50 μm.
j. Corresponding quantification of KLF4 nuclear intensity (normalized to the average of 2i value in corresponding experiment). Averaged data from 3 independent experiments (n = 41/44/43/35/29/18/30 for 2i-0hr/2i-24hr/N2B27-24hr/2i-48hr/N2B27-48hr/2i-72hr/N2B27-72hr). The effect of media is significant (Two-way ANOVA).
k. Corresponding quantification of KLF4 nuclear intensity in 2i, colony edge vs interior (n = 41/44/35/18/ for 2i-0hr/2i-24hr/2i-48hr/2i-72hr). Averaged data from 3 independent experiments. The effect of edge vs interior is not significant (Two-way ANOVA with repeated measures).
l. Corresponding quantification of KLF4 nuclear intensity in N2B27, colony edge vs interior (41/43/29/30 for 2i-0hr/N2B27-24hr/N2B27-48hr/N2B27-72hr). Averaged data from 3 independent experiments. The effect of edge vs interior is not significant (Two-way ANOVA with repeated measures).

Error bars show mean ± s.e.m.

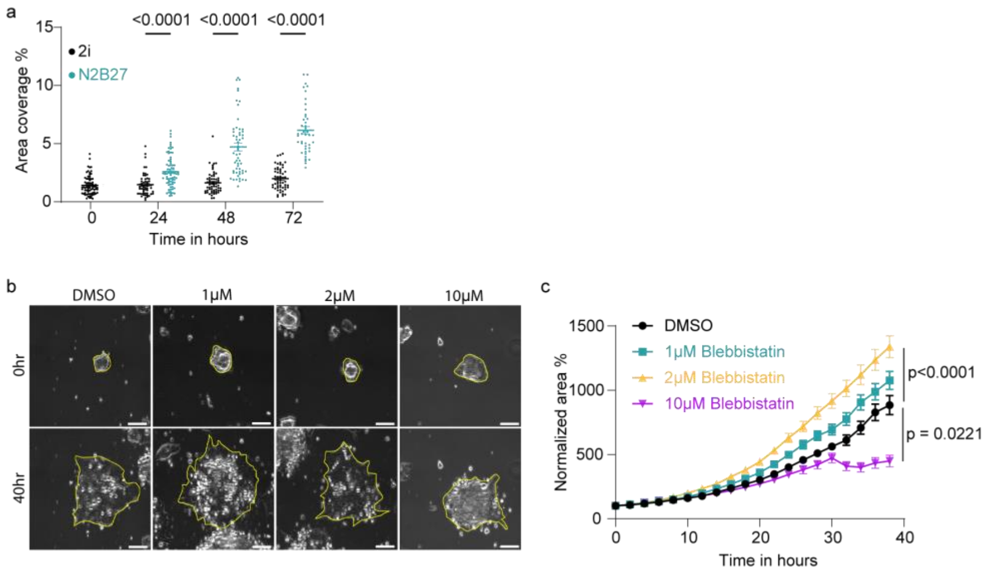


**Supplementary Figure 2. Further characterization of focal adhesions and effects in colony spreading area caused by blebbistatin treatment.**

a. Quantification of the density of phospho-paxillin adhesions. Data is from 3 independent experiments (n = 66/56/73/58/56/56/44 colonies for 2i-0hr/2i-24hr/N2B27-24hr/2i-48hr/N2B27-48hr/2i-72hr/N2B27-72hr). The effect of media and time is significant (Two-way ANOVA).
b. Representative phase contrast images of mESC colonies grown in indicated concentrations of blebbistatin for 40 hours. Colony borders are demarcated in yellow. The scale bar is 50 μm.
c. Quantification of colony area evolution (normalized to initial value) with time for mESC colonies grown in mentioned media. Data from 3 independent experiments (n=70/75/66/39 for DMSO/1μM-/2μM-/10μM-Blebbistatin treatments, Kruskal-Wallis test, comparisons with respect to DMSO for the last time point).

Error bars show mean ± s.e.m.

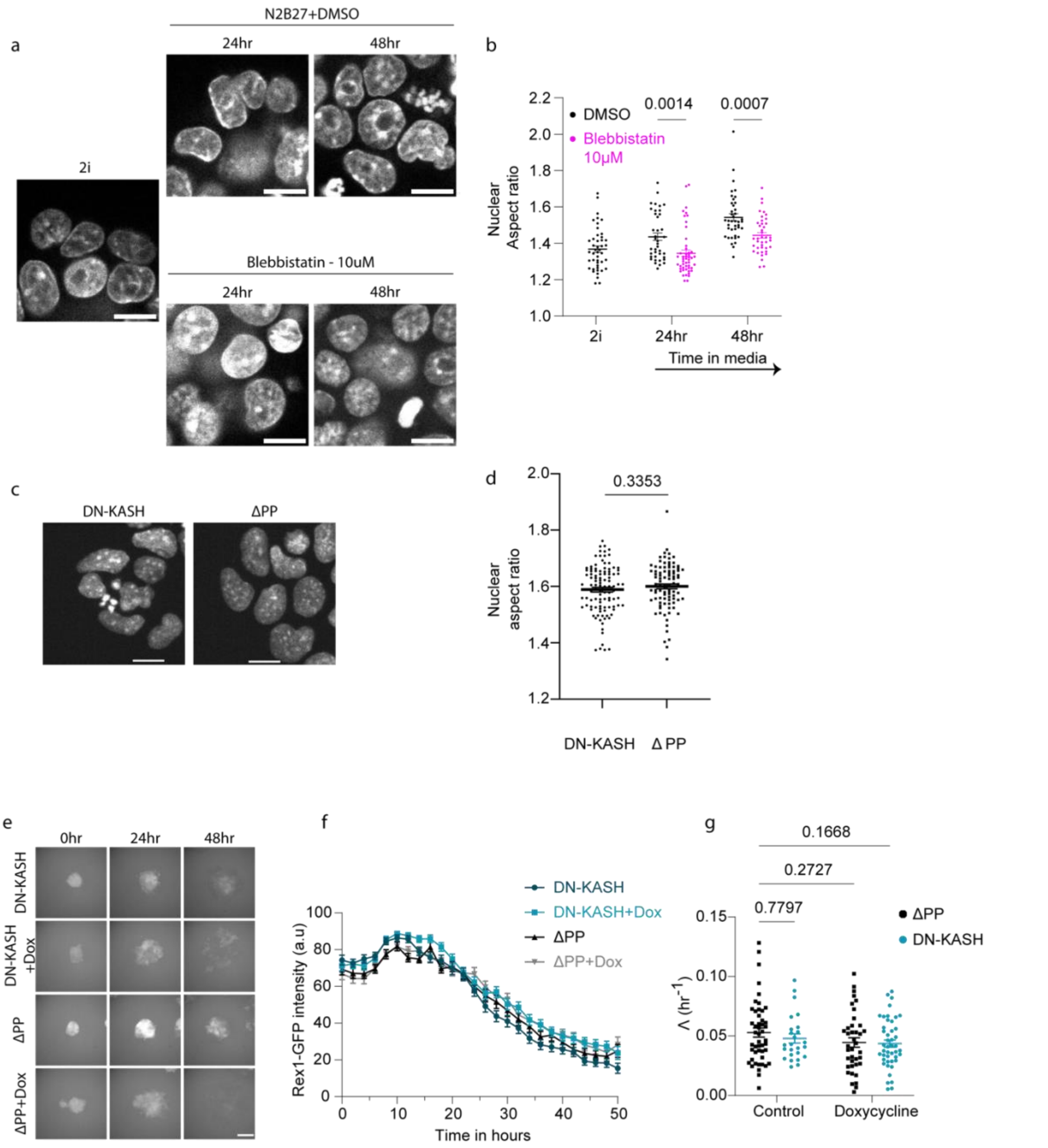


**Supplementary figure 3. Further quantifications on nuclear shapes and DN-KASH overexpression.**

a. Representative image of mESC colonies grown in 2i/N2B27 supplemented with DMSO or 10µM blebbistatin for mentioned time points and stained for nucleus with Hoechst. The scale bar is 10µm.
b. Quantification of Nuclear aspect ratio averaged over colony area. Averaged data from 3 independent experiments (n = 45/39/50/41/40 colonies for 2i-0hr/N2B27+DMSO-24hr/N2B27+ 10µM-Blebbistatin –24hr/N2B27+DMSO-48hr/N2B27+10µM-Blebbistatin –48hr). The effect of treatment is statistically significant (Two-way ANOVA without repeated measurements).
c. Representative fluorescence images of mESC colonies of DN-KASH/ΔPP expressing mESCs for the corresponding time points in N2B27 on 5kPa polyacrylamide gels. Doxycycline was added at 0hr (along with N2B27). The scale bar is 10µm.

d. Quantification of Nuclear aspect ratio averaged over single mESC colonies. Averaged data from 4 independent experiments (n = 105/92 for DN-KASH+Doxycycline/ΔPP+Doxycycline). The effect is not significant (Unpaired t-test).
e. Representative fluorescence images of mESC colonies of DN-KASH/ΔPP genetic background grown for mentioned time points in N2B27 on 5kPa polyacrylamide gels. Doxycycline and N2B27 were added at 0hr. The scale bar is 100µm.
f. Corresponding quantification of normalized Rex1-GFP intensity, as a function of time for each genetic background. Averaged data from 3 independent experiments (n = 25/48/45/39 colonies for DN-KASH/DN-KASH+Doxycycline/ΔPP/ΔPP+Doxycycline).
g. Quantification of Rex1-GFP decay constants. Data from 3 independent experiments (n = 25/48/45/39 colonies for DN-KASH/DN-KASH+Doxycycline/ΔPP/ΔPP+Doxycycline). The effect of DN-KASH is significant only upon Doxycycline induction (Two-way AOVA without repeated measurements).

Error bars show mean ± s.e.m.

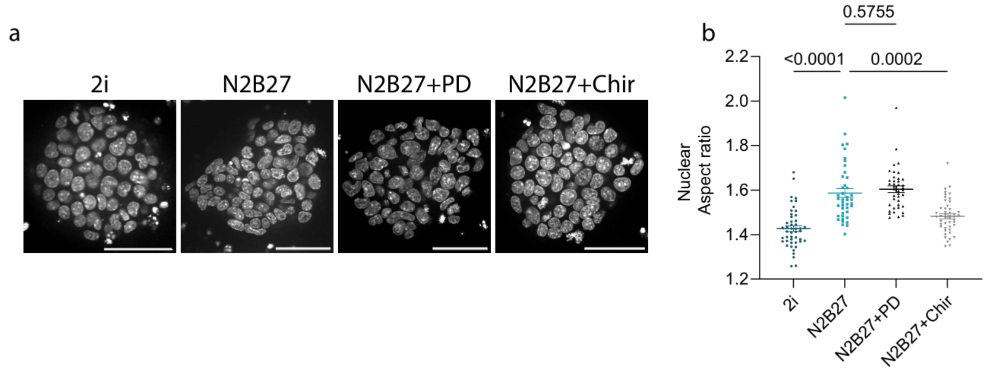


**Supplementary Figure 4. Nuclear aspect ratios upon single inhibitor treatments.**

a. Representative images of mESC colonies grown in the corresponding media and stained with nucleus marker Hoecsht. The scale bar is 50µm.
b. Quantification of nuclear aspect ratio averaged for mESC colonies grown in mentioned media for 48 hours. Averaged data from 3 independent experiments (n=47/42/40/47 colonies for 2i/N2B27/N2B27+PD/N2B27+Chir). The effect of media change is not significant only for the comparison between N2B27 and N2B27+PD (Kruskal-Wallis test, comparisons with N2B27)

Error bars show mean ± s.e.m.

**Supplementary video legends**

**Supplementary video 1.** Time-lapse of mESC cells seeded on 2i versus N2B27 conditions. Example of mESC colonies showing Rex1-GFP signal (left), phase contrast images (center) and traction forces (right). Top images, 2i conditions, bottom images, N2B27 conditions. The scale bar is 20 µm. Values of the heatmap represent traction forces in pascals.

**Supplementary video 2.** Time-lapse of mESC cells with or without blebbistatin. Example of mESC colonies in N2B27 medium showing Rex1-GFP signal (left), phase contrast images (center) and traction forces (right). Top images, control conditions (DMSO), bottom images, 2 µm blebbistatin. Scale bar is 20 µm. Values of the heatmap represent traction forces in pascals.

**Supplementary video 3.** Time lapse of mESC cells with/without GSK3 or ERK signaling inhibition. Example of mESC colonies in different media combinations showing Rex1-GFP signal (left), phase contrast images (center) and traction forces (right). Top images 2i condition, second row N2B27, third row N2B27 with ERK signaling nhibitor PD0325901 (N2B27+PD) and bottom row N2B27 with GSK3 agonist CHIR99021 (N2B27+Chir). The scale bar is 20 µm. Values of the heatmap represents traction forces in pascals.
